# Comparative analysis of genome-wide protein-DNA interactions across domains of life reveals unique binding patterns for hypersaline archaeal histones

**DOI:** 10.1101/2022.03.22.485428

**Authors:** Saaz Sakrikar, Rylee K. Hackley, Mar Martinez-Pastor, Cynthia L. Darnell, Angie Vreugdenhil, Amy K. Schmid

**Affiliations:** Biology Department, Duke University, Durham, NC, USA; University Program in Genetics and Genomics, Duke University, Durham, NC, USA; Center for Computational Biology and Genomics, Duke University, Durham, NC, USA

## Abstract

DNA-binding proteins with roles in chromatin architecture and transcriptional regulation are present in all three domains of life. Histones package DNA and regulate gene expression in eukaryotes, and find their evolutionary origin in the domain of life Archaea. Previously characterised archaeal histones have a somewhat conserved functional role in nucleosome formation and DNA packaging. However, previous research has indicated that the histone-like proteins of high salt-adapted archaea, or halophiles, appear to function differently. The sole histone protein encoded by the model halophilic species *Halobacterium salinarum* is non-essential, is involved in direct and indirect transcriptional regulation, and does not appear to package DNA. Here we use protein-DNA binding assays, computational analysis, and quantitative phenotyping to compare DNA binding patterns across halophilic histone proteins, bacterial and archaeal TFs, NAPs, and eukaryotic histones. Like TFs, halophilic histones bind the genome too sparsely to compact the genome. However, unlike TFs, binding occurs in both coding and intergenic regions. Unlike histones, halophilic histone occupancy is not depleted at the start sites of genes, and halophilic genomes lack the dinucleotide periodicity known to facilitate histone binding. We detect unique sequence preferences for histone binding in halophiles. Together these data suggest that the non-essentiality and genome-wide binding features of halophilic histone-like proteins are conserved across halophiles; they bind DNA in ways resembling both TFs and chromatin proteins, but do not appear to play a role in forming chromatin.

**IMPORTANCE:** Most cells in eukaryotic species – from yeast to humans– possess histone proteins that pack and unpack DNA in response to environmental cues. These essential proteins regulate the genes necessary for important cellular processes, including development and stress protection. The domain of life Archaea represent the evolutionary progenitors of eukaryotes. The universal conservation of the primary sequences of histone proteins across archaeal lineages suggests that eukaryotic histones originated in the Archaea. However, archaeal histones lack N-terminal tails and, in some species, package DNA in a continuous helix with no linker DNA between nucleosomes. We recently discovered that histones in hypersaline adapted archaeal species do not package DNA, and can act like transcription factors (TFs) to regulate stress response gene expression. Here we compare hypersaline histone function to a variety of DNA binding proteins across the tree of life, revealing a mosaic of functions for hypersaline-adapted histones.

## INTRODUCTION

Formation of chromatin and regulation of transcription are fundamental features shared by all domains of life and are mediated by a variety of DNA binding proteins. These proteins are commonly characterised as chromatin proteins(1) and transcription factors (TFs)(2–4) respectively, with some known to play both architectural and gene regulatory roles(5). The amino acid sequences of many of these proteins are conserved throughout the domains of life. Histones are ubiquitous in eukaryotes, where their architectural role is well-characterised(6), and are also encoded in genomes across most archaeal lineages(7). Transcription factors with helix-turn-helix DNA binding domains are widespread in all domains of life(4, 8, 9). In contrast, some chromatin proteins are specific to certain clades within the domain Archaea, for example Mc1(10) and Cren7(11).

However, conserved primary sequence and even protein structure does not necessarily imply conserved function. For instance, Alba, which can bind both DNA and RNA, is involved in chromatin architecture and in RNA structure stability in Archaea, while in eukaryotes Alba acts in greater variety of pathways, particularly in translation(12). While Alba in both domains is capable of binding DNA and RNA, its specific biological roles have diverged both within and across the domains of life. Similarly, the histone fold domain is present in all archaeal histones as well as eukaryotic core histones(13), but is also detected in some eukaryotic transcription factors(14). Again, while the DNA-binding role of the histone fold is conserved, its precise cellular function has diverged.

In understanding the function of these DNA-binding proteins, the question of which features of molecular function are conserved remains open: how do these proteins play architectural roles, how do they regulate transcription, or perform both functions? Certain hallmarks give clues as to the cellular functions of such proteins, including sequence determinants of binding, position relative to genomic features (i.e., coding or noncoding DNA), size of binding footprints, and shape of binding peaks. Many of these features are detectable in genome-wide DNA binding location data such as ChIP-seq (chromatin immunoprecipitation coupled to sequencing)(15–19). Here, we investigate the role of the histone-fold containing proteins found in halophilic archaea with phenotypic analysis and ChIP-Seq, and by comparing it with ChIP-Seq data from other DNA-binding proteins to establish its binding characteristics based on the hallmarks described above.

Across bacterial species, nucleoid-associated proteins (NAPs) share architectural roles in DNA but differ in their mechanism of binding(20–22). Some NAPs such as HU(21) or Lrp(23) bind without sequence specificity, while others (like IHF(21)) bind a defined cis-regulatory sequence motif. NAPs are also diverse in terms of genomic binding locations: HU binds throughout the genome without discrimination between coding and non-coding regions(21), whereas H-NS and Fis preferentially bind promoter regions(20). Regardless of precise genomic location, NAPs bind frequently, covering 10-20% of the genome(20, 21). This is consistent with high NAP protein expression levels(24), and their functions in genomic structural organization(22). NAPs typically affect transcription globally, including indirect effects far from their binding loci(21, 25).

In archaeal transcription, proteins that guide RNA polymerase to core promoters resemble those of eukaryotes (TATA binding protein, transcription factor B), whereas TFs that regulate gene expression in response to environmental perturbation more closely resemble those of bacteria(4). Haloarchaeal and bacterial TFs typically regulate transcription of target genes by binding in a sequence-specific manner(26, 27) proximal to gene promoters(4). These proteins regulate expression of target genes by recruiting or hindering the basal transcriptional machinery(4). However, in bacterial and archaeal species, some TFs bind hundreds of sites in the genome and can bend or loop DNA, such as Lrp family homologs(15, 28). DNA binding specificity is often weak when such TFs play architectural roles(15, 23, 28, 29).

In eukaryotes, the histone fold domain is required in the formation of the histone octamer (nucleosome core) and for histone-DNA binding(30). Eukaryotic nucleosomes package DNA and regulate gene expression(6). Like bacterial NAPs, eukaryotic core histones play an important role in genome architecture, binding frequently throughout the genome. Histone binding to DNA leads to nucleosome formation, but active promoter regions form “nucleosome-depleted regions” due to reduced histone binding, while regularly spaced nucleosomes are present downstream of the gene start(16, 31, 32). While these histones do not have a defined sequence motif, they disfavour poly-A sequences and preferentially bind sequences with ~10bp periodicity of A/T dinucleotides(16, 32, 33). This corresponds with the length of the helical pitch of DNA(34), and the positioning of the A/T dinucleotides is hypothesized to aid wrapping of the DNA helix around the nucleosome(33). Like NAPs, histones are expressed at very high levels in eukaryotes (and in archaeal species in which histones play architectural roles) to bind and compact the genome(35–37).

Phylogenetic evidence has traced the origin of the histone fold domain to the archaeal domain of life(13, 38, 39). Studies primarily in thermophilic archaea representing the euryarchaeal superphylum suggested a eukaryotic-like DNA packaging function of these archaeal histones(40–42), which can form nucleosomes (like in eukaryotes). Extended polymeric structures known as hypernucleosomes have also been observed that wrap and compact the genome in multiples of 30-60 bp(42–44). The genomes of many of these well-studied species encode two histone paralogs; deletion of a single histone gene was found to be viable, while a deletion of both was not possible(45). These archaeal histones favour binding sites with 10-bp periodicity of AT-dinucleotides(46), similar to eukaryotic histones(33). Indeed, a genome-wide 10bp periodicity signal was detected in several archaeal species with histones(47). In addition to histones, a number of non-histone proteins of bacterial origin as well as archaeal-specific proteins also contribute to DNA architecture in the Archaea(7). For example, most Crenarchaeaota lineages lack histone fold proteins, instead utilizing the NAPs Alba, Cren7, and Sul7d for genome compaction(12, 48). *Thermoplasma acidophilum* also does not encode histones. Instead the role of genome compaction is performed by a homolog of the bacterial-like NAP, HU(49). In species of *Methanosarcina*, the histone protein was found to be dispensable for growth, and an archaeal protein, Mc1, was shown to be capable of performing an architectural role in the genome(10, 50). Hence, while archaeal histones perform their conserved role in chromatin architecture in many thermophilic species of archaea, other NAPs are also involved in archaeal genome architecture, and the role of histones in non-thermophilic lineages remains unclear.

In the hypersaline adapted (halophilic) lineage of archaea, the cellular function and DNA binding properties of histone-like proteins are understudied. So far, evidence points to a specialized structure and alternative function for halophilic histones. Halophilic archaeal genomes encode a sole histone protein with two histone fold domains(51, 52). This fused heterodimer forms a nucleosome with a structure resembling the (H3-H4) dimers present at the core of eukaryotic nucleosomes, but with a long flexible linker between monomers(51) (Schmid, unpublished data from RosettaFold predictions(53)). This fused dimer is strongly conserved across halophile genomes(52). As an adaptation to their hypersaline environment, halophilic archaea have evolved a “salt-in” strategy by which the external sodium concentration is balanced by a very high (~3-4 M KCl) potassium concentration within the cytoplasm(54). The resultant highly ionic cytoplasm has led to further adaptations: the halophilic proteome is highly acidic to aid stability and solubility in this charged environment(54). Hence, the surface of halophilic histones is highly acidic, unlike known eukaryotic histones in which the basic surface facilitates electrostatic attraction to DNA(52, 55).

Our previous work demonstrated that putative DNA binding proteins encoded in *Halobacterium salinarum*, including the histone-like protein HpyA and putative NAPs, are expressed at levels too low to be able to compact the genome(52). HpyA binding is sparse throughout the genome (~60 sites(56)) and it can be deleted with no growth defect under standard conditions(52).

Instead, HpyA functions as a transcriptional regulator of inorganic ion transport and nucleotide metabolism by binding DNA in low sodium conditions(56). HpyA is necessary for growth and cell shape maintenance in low salt, therefore linking its regulatory function to hypo-osmotic stress resilience(56). Although HpyA cellular functions have been investigated, its DNA binding properties and conservation of functional features with other DNA binding proteins across the tree of life remain unclear.

To reveal the true evolutionary trajectory of DNA binding proteins that integrate genomic architecture with transcription regulation, a more thorough understanding of archaeal histone function across the domain is required. Here we ask how and whether halophile histone binding properties are conserved across halophiles and the domains of life. The binding characteristics of these halophilic histone proteins are compared to those of characterised bacterial NAPs, archaeal and bacterial TFs, and archaeal and eukaryotic histones. We used quantitative phenotyping data, ChIP-seq, and genomic sequence data to evaluate halophilic histone function and binding attributes based on six criteria: (i) knockout phenotype; (ii) sequence specificity; (iii) binding location (intergenic vs coding); (iv) binding frequency; and (v) binding peak size and shape. We examine the halophilic histone proteins from two model halophilic species, *Hbt. salinarum* and *Haloferax volcanii*. These species represent two different clades of the halophilic phylogeny and are therefore representative models for halophiles in general(57). Our results suggest that haloarchaeal histone proteins are largely conserved across halophilic species and possess functional attributes that differ from other known DNA binding proteins in some respects but are similar in others. Halophilic histones therefore blur the distinctions between TFs, nucleoid proteins, and histones, calling for more nuanced definitions of DNA binding proteins.

## RESULTS and DISCUSSION

### *Haloferax volcanii* histone HstA is not essential but important for maintaining wild type growth rate

HstA (HVO_0520) is the sole histone protein encoded in the genome of the model halophile *Hfx. volcanii*. As we observed in our previous study(52), HstA shares 65% sequence identity with HpyA histone-like protein of *Hbt. salinarum* and retains residues conserved across histones of nearly 80 sequenced halophile genomes. Here we used a genetic approach to determine if our previous observations regarding non-essentiality of the *Hbt. salinarum* histone HpyA are generalizable to other species of halophiles(52, 56). We observed that *hstA* was readily deleted from *Hfx. volcanii* (details in Materials and Methods); however, unlike the Δ*hpyA* deletion strain of *Hbt. salinarum*, this Δ*hstA* strain exhibited a slight but significant growth defect compared to the parent strain in optimal conditions (rich media at 42°C, **Fig. 1A and B** and **Table S1**, https://doi.org/10.6084/m9.figshare.19391648.v1. 84% of parent strain growth as measured by area under the curve; Welch two-sample t-test *p* < 1.5 x 10^−6^). Growth under a variety of stress conditions (sodium and magnesium stress, oxidative stress with peroxide, alternate nutrient conditions) was tested. The ratio of the area under the curve for Δ*hstA* to parent under these conditions was found to be in the range 84%-91%, at or above the ratio for optimal conditions (**Fig. S1**). These data indicate that the Δ*hstA* growth defect under standard conditions is not further compounded under these stress conditions, suggesting that *hstA* is dispensable for growth under the stress conditions tested. This growth defect is significantly complemented by the *in trans* expression of *hstA* alone or translationally fused in frame to the hemagglutinin epitope tag (**Fig. S2**), indicating that the growth defect is attributable to the deletion of *hstA* and not due to polar effects on surrounding genes. In addition, these data indicate that the C-terminal HA tag does not interfere with HstA function, allowing ChIP-seq with the tagged strain to be carried out. Whole-genome re-sequencing verified the absence of any secondary site mutations in this strain (**Table S1**) and the complete absence of any wild type *hstA* copies from the genome (halophiles are highly polyploid(58), necessitating such validation). Taken together, these results establish that *hstA* is a non-essential gene whose deletion causes a slight but significant growth defect under standard growth conditions. We conclude that HstA resembles *Hbt. salinarum* HpyA in that both are dispensable for growth. In contrast, known chromatin proteins are usually essential(59, 60), if not individually, then combinatorially. For example, in *Thermococcus kodakarensis*, with two histone genes, either single knockout is viable but a double knockout is not possible(45). HpyA and HstA therefore differ from other known histones with respect to their essentiality for viability.

**Figure 1:**
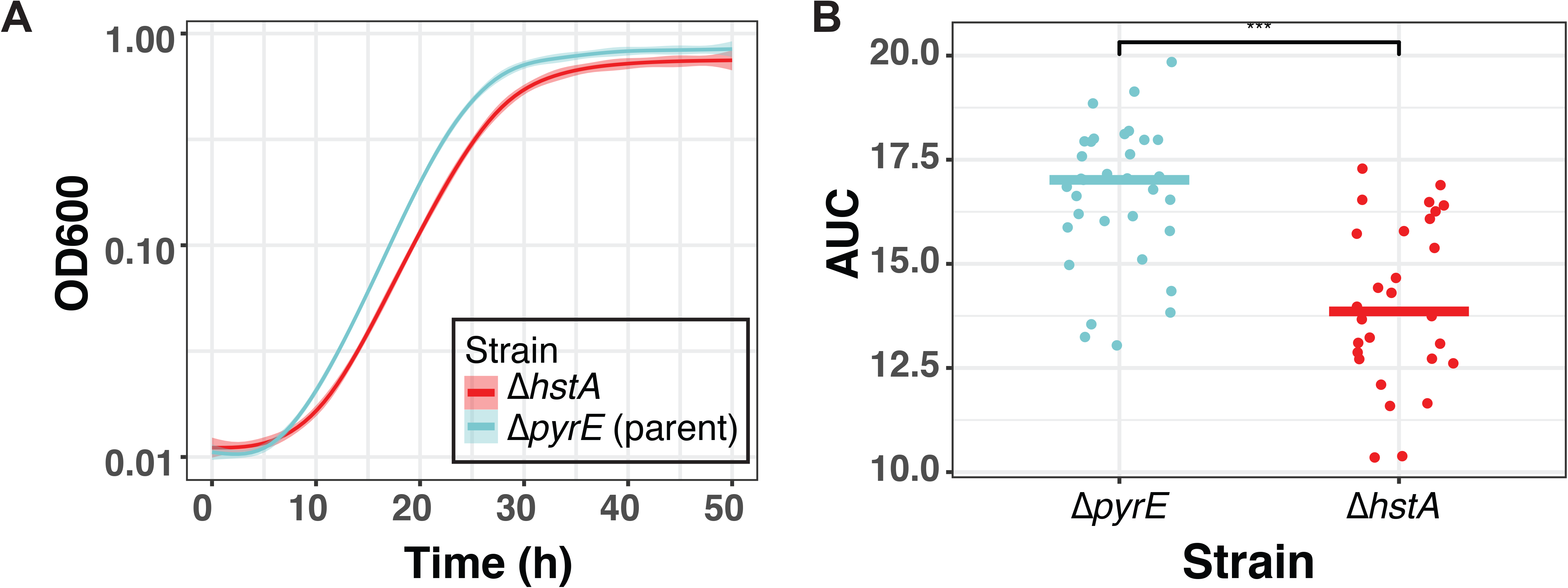
Δ*hstA* strain is impaired for growth in optimal conditions. (**A**) Growth of strains (Δ*pyrE* blue, Δ*hstA* red; 9 biological replicates with 2-3 technical replicates each) measured as optical density (OD600). The heavy lines represent the smoothed conditional mean growth curves, shaded area surrounding each curve represents the error of the mean. (**B**) Area under the curve (AUC) of the growth curve for each strain under standard conditions as calculated by R package grofit; each dot represents AUC for one technical replicate growth curve. Horizontal lines represent the median of the distribution of points for each strain.

### HstA genome-wide location analysis (ChIP-seq) reveals binding patterns like those of HpyA

We carried out chromatin immunoprecipitation coupled to sequencing (ChIP-seq) to locate and compare genome-wide binding sites of HstA to HpyA(56) (details in Materials and Methods). Each ChIP-seq experiment was carried out under conditions in which each halophilic histone would be expected to be active in DNA binding. Given that HstA is important from growth under standard conditions, we conducted ChIP-seq experiments in optimal conditions in exponential phase, whereas HpyA is active (and ChIP-seq conducted) in low salt conditions. Across the *Hfx. volcanii* genome, we observed infrequent HstA binding in discrete, sharp peaks of enrichment (**Fig. 2A**). This binding frequency and peak shape were like those observed for *Hbt. salinarum* HpyA(56) (**Fig. 2B)**. Binding peak shape in ChIP-seq data correlates with function: such sparse, punctuated narrow peak shapes are often observed for site-specific TFs, in contrast to the wide, broad peaks typical of histones(17). The similarities between the binding profiles of the two histone-like proteins extend further: the number of reproducible peaks for HstA was 32 (**Table S2**), on the same order of magnitude seen for HpyA (59 peaks(56)). HstA peaks averaged 374 bp wide (see zoom-in of representative peak, **Fig. 2A**), the same order of magnitude as the mean peak width observed for HpyA (299 bp, **Fig. 2B, Table S3**). For each of HpyA and HstA, these binding loci cover <1% of the genome of *Hbt. salinarum* and *Hfx. volcanii*, respectively, suggesting that the overwhelming majority of the genome is not bound by halophilic histones. Of these HstA peaks,15.6% were in non-coding regions, which closely corresponds with the 15.8% of the genome that is intergenic; hence, like HpyA(56), HstA binding is not enriched in non-coding regions (*p*-value >0.4; **Table S4**). However, unlike HpyA, which regulates ion uptake(56), the genes nearby HstA binding peaks were not enriched for a particular function according to archaeal clusters of orthologous genes (arCOG) categories(61). Taken together, these data suggest that the genome-wide DNA binding patterns of *Hbt. salinarum* and *Hfx. volcanii* histone-like proteins are conserved across halophiles, with their binding peak frequency and shapes resembling those of TFs.

**Figure 2:**
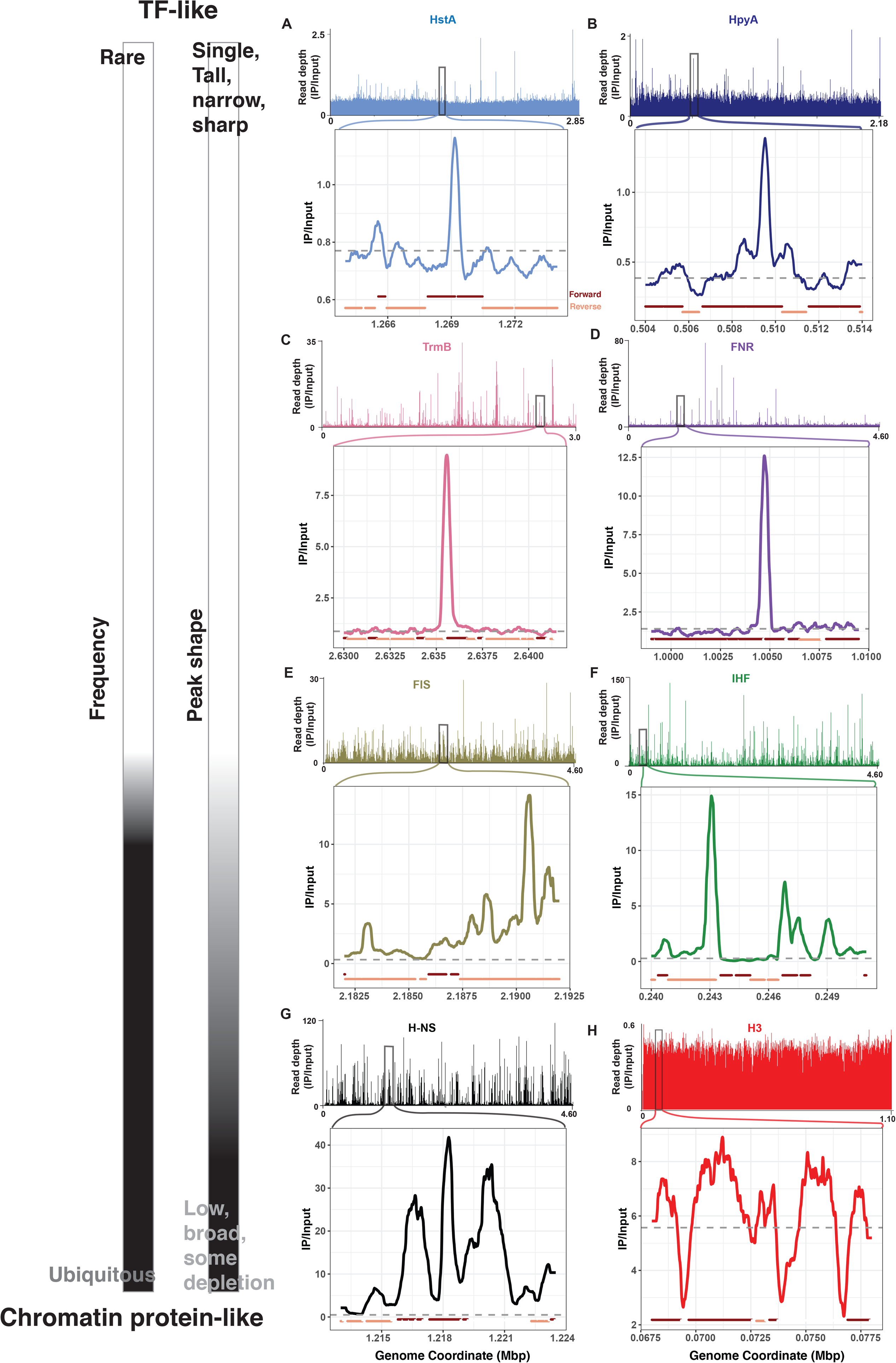
ChIP-seq binding signal for HpyA and HstA compared with TFs, NAPs, and eukaryotic histone. In each panel, chromosome-wide binding patterns (measured as read-depth of IP/Input) are shown above and zoomed-in regions of representative peaks are shown below. All archaeal and bacterial genome views depict the main chromosome of each species. (**A and B**) show halophilic histone-like protein binding patterns: (**A**) *Hfx. volcanii* HstA (light blue, NCBI accession NC_013967.1, peak centre 1.27Mb); (**B**) *Hbt. salinarum* HpyA (dark blue, NC_002607.1, peak centre 0.51Mb). (**C**) Depicts halophilic TF *Haloarcula hispanica* TrmB (pink, NC_015948.1, peak centre 2.64Mb). (**D**) Shows bacterial TF *E. coli* FNR (purple, NC_000913.3, peak centre 1.01Mb). (**E-G**) *E. coli* NAPs: (**E**) H-NS (black, peak centre 1.22Mb); (**F**) IHF (green); (**G**) Fis (olive). (**H**) Shows yeast histone H3 (red), chromosome VII (NC_001139.9). For the TFs and H-NS, known to directly regulate target genes(20, 26, 62), peaks with a known functional role were chosen. For each genome-wide view and zoom-in, X-axis represents chromosomal coordinates in Mbp, Y-axis represents the read depth ratio of IP to input control (i.e., binding enrichment). Grey dotted line in the zoom-ins represent a baseline calculated from the average genome-wide IP/Input signal; dark red and tan lines below each zoom-in plot represent genomic context (forward and reverse strand genes, respectively). Scale at left indicates the classification of each DNA binding protein pattern based on features of frequency and peak shape.

### Comparison of halophilic histone-like protein binding patterns with those for TFs, NAPs, and eukaryotic histones

To understand the DNA binding functions of halophilic histones relative to known DNA-binding proteins, we compared our ChIP-seq results to those of eukaryotic histones, bacterial NAPs and TFs, and halophilic TFs (details of datasets used in Methods section and **Table S5**). Together, these proteins encompass a wide spectrum of cellular functions and DNA binding modes, and hence provide a comprehensive set of comparisons for our data from halophilic histones.

We first analysed the similarities and differences between these proteins by visual inspection of the genome-wide binding patterns and representative zoomed-in regions (**Fig. 2, Fig. S3)**. We focused on peak shape and genome-wide binding frequency: it has been observed that “canonical” TFs tend to exhibit tall, sharp peaks and bind rarely(17), in concordance with their role in site-specific regulation of transcription initiation(4). As described above, we observed that halophilic histone-like proteins exhibit discrete, narrow regions of binding enrichment at relatively few locations in the genome (**Fig. 2A, B, S3A)**. We observe similarly sharp, discrete peaks binding rarely in the case of halophilic TF TrmB (**Fig. 2C, Fig. S3B)** and bacterial TF FNR(62) (**Fig. 2D, Fig. S3C)**. Such punctuated peak shapes have been observed for sequence-specific TFs in eukaryotes as well(17). In contrast, previous research has shown chromatin-like proteins (histones, NAPs) bind ubiquitously with broader, flatter peaks, and/or with areas of depletion, in keeping with their roles in DNA architecture and compaction(17, 19, 21, 29). Consistent with this, we observed that bacterial NAPs IHF, H-NS, and FIS bound frequently in the genome (**Figs 2E-G, Fig. S3D**), particularly for H-NS and IHF, corroborating previous reports that binding sites of these NAPs cover a significant portion of the genome(20). However, we observed a mixture of peak shapes for the NAPs. H-NS exhibited very broad peaks, often more than a kb in width (**Fig. 2G**), while there was mix of narrow and broad peaks seen for IHF (**Fig. 2F**) and FIS (**Fig. 2E**), with a very high frequency of peaks for IHF (also seen in **Fig. S3D**). Indeed, for yeast histones, we do not observe binding peaks at all in the genome-wide view (**Fig. 2H, Fig. S3E**). Upon closer inspection of local genomic regions, we observe broad, flat areas of enrichment punctuated by depletion at promoter regions, as expected from ubiquitous binding and nucleosome formation (except at promoters of transcribed genes, **Fig. S3E**)(16, 32).

To quantify these characteristics to gain further insight into halophilic histone function, we tallied and compared the number of binding events, average peak width, and percentage of the genome covered for each DNA binding protein (Methods, **Table S3**, **Fig. 3)**. As discussed above, we detected 59 reproducible binding sites for HpyA(56) and 32 for HstA. This is at the lower end but within the range observed for haloarchaeal and bacterial TFs (36-253 peaks, **Table S3**). The average width of HpyA and HstA peaks (299bp and 374bp, respectively) is comparable to the range of peak widths observed for haloarchaeal TFs (318-466bp; **Fig. 3A**) and the bacterial TF FNR (511 bp average width). Additionally, halophilic histone peaks most closely resemble halophilic and bacterial TFs with respect to the percentage of the genome bound (<1% for HpyA and HstA, 0.4-2.2% for haloarchaeal TFs, 2% for bacterial TF; **Table S3**, **Fig. 3B**). By contrast, the average peak width and genome coverage is more variable across the various bacterial NAPs included in the comparison here (**Fig, 3A, B**). On average, NAP binding sites are more numerous (mean number of peaks is 631), wider (average 1.2kb), and cover more of the genome (average 14%) than HpyA and HstA binding peaks, particularly in the case of H-NS (**Table S3**, **Fig. 3**). These numbers generated from our analysis correspond with published estimates that, because of their genome-wide architectural role, NAPs cover 10-20% of the genome(20, 21).

**Figure 3:**
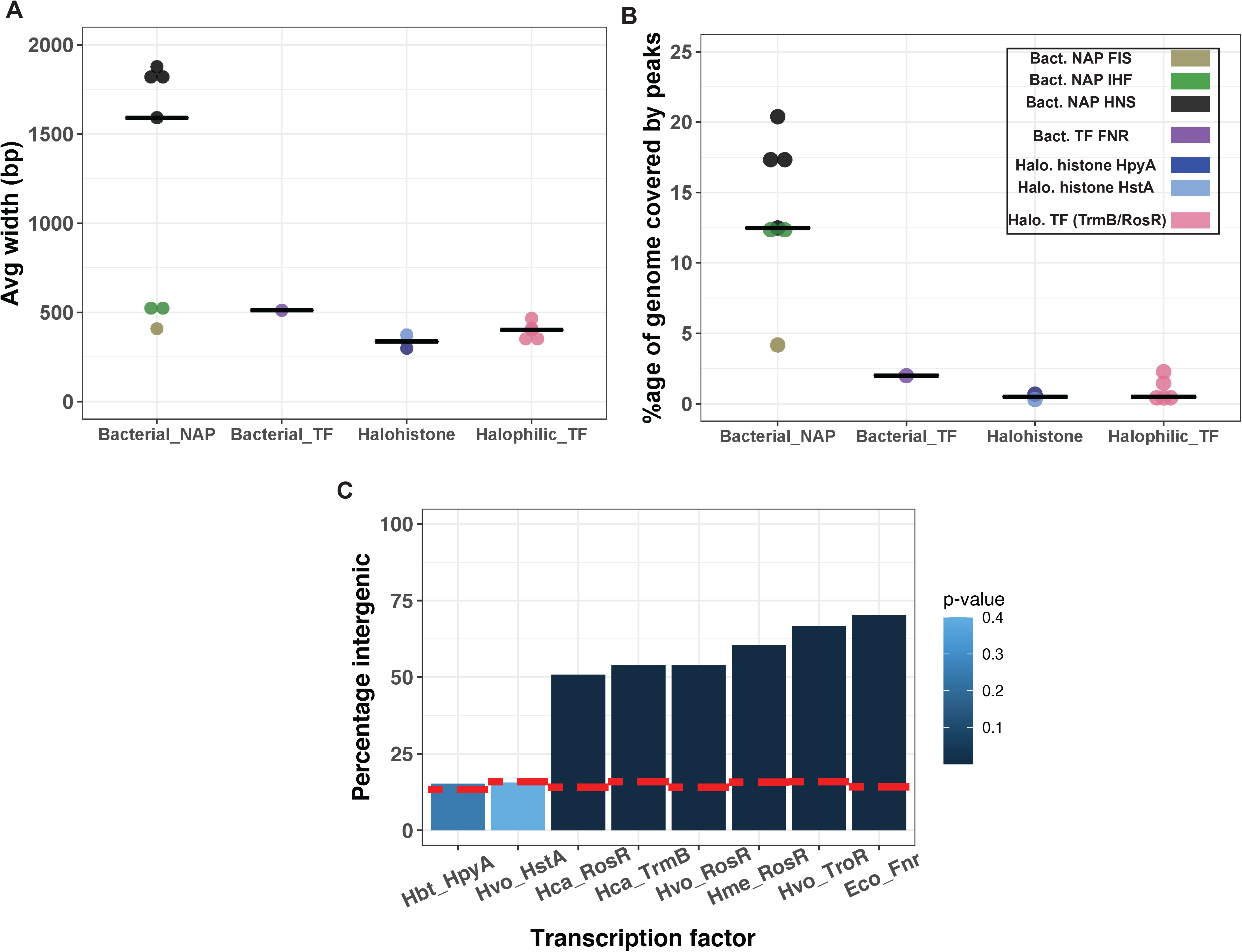
Genomic features of HpyA and HstA binding sites according to ChIP-seq data. (**A**) Average width of all ChIP-seq peaks for a given DNA-binding protein, arranged into columns by type (bacterial NAPs, bacterial TFs, halophilic histones, halophilic TFs). (**B**) Percentage of genome covered by all ChIP-seq peaks of a given DNA-binding protein arranged into columns by type (bacterial NAPs, bacterial TFs, halophilic histones, halophilic TFs). (**C**) Bar graph of ChIP-seq peaks for HpyA, HstA, and transcription factors TrmB, RosR, and TroR. Species names are abbreviated: Hbt, *Hbt. salinarum;* Hvo, *Hfx. volcanii*; Hca, *Hca. hispanica;* Eco, *E. coli*. The height of each bar represents the percentage of peaks located in intergenic regions. Dotted red line indicates the percentage of each genome that is non-coding. The intensity of colour of the bars represents hypergeometric test *p*-values of significance for enrichment within promoter regions (see legend for colour scale).

Because binding location often relates to molecular function, we next analysed the location of the HpyA and HstA peaks relative to genomic features (**Fig. 3C)**. As discussed above, neither HpyA nor HstA show a preference for certain genomic features, neither coding nor promoter sequences. In contrast, as expected from previous studies(26), haloarchaeal and bacterial TF binding sites were significantly overrepresented in intergenic regions relative to the genomic backgrounds of these species, which are 84-87% coding (hypergeometric test; p<10^−3^) (**Table S4)**. For these TFs, the proportion of peaks binding to intergenic regions varied between 51% for *Hfx. mediterranei* TrmB to 70% for *E. coli* FNR (**Fig. 3C**). Hence, while halophilic histones appear to bind without preference for genic or intergenic regions, TF binding favours intergenic regions.

Taken together, these data demonstrate that halophilic histones meet many qualitative and quantitative criteria for binding patterns like those of TFs (binding at discrete, narrow peaks at relatively few genomic sites). However, in their lack of preference for binding particular genomic features, HpyA and HstA resemble NAPs such as IHF and HU(21). Thus, HpyA and HstA binding patterns resemble those of TFs in some respects but non-specific DNA binding proteins in others.

### Haloarchaeal histone-like protein occupancy curves surrounding start sites are unique relative to canonical histone and TF signals

To further challenge the hypothesis that HpyA and HstA bind DNA like typical histones, the average occupancy (normalized read depth) at open reading frame (ORF) start sites were compared across DNA binding proteins (see Methods). As expected from previous studies of global nucleosome occupancy,(16, 31, 63, 64) we detected a depletion in ChIP-seq binding signal at promoter regions but enriched at regularly spaced nucleosomes in the gene body for *Saccharomyces cerevisiae* histones H3, H4, and H2B **(Fig. 4A**, data sources listed in **Table S5)**. Specifically, an upstream decline in occupancy was detected, with a minimum at ~160-200 bp upstream of the start site for all histone proteins analysed. In contrast, binding peaks representing the +1 to +3 nucleosomes bound downstream of the start site were detected in the expected positions for histone H3 (+1 at ~100 bp, with the subsequent peaks at ~150 bp intervals, **Fig. 4A**). The nucleosome positions detected in our analysis therefore correspond with known positions of the nucleosome-free region and bound nucleosomes across gene start sites and into the 5’ end of gene coding regions(16).

**Figure 4:**
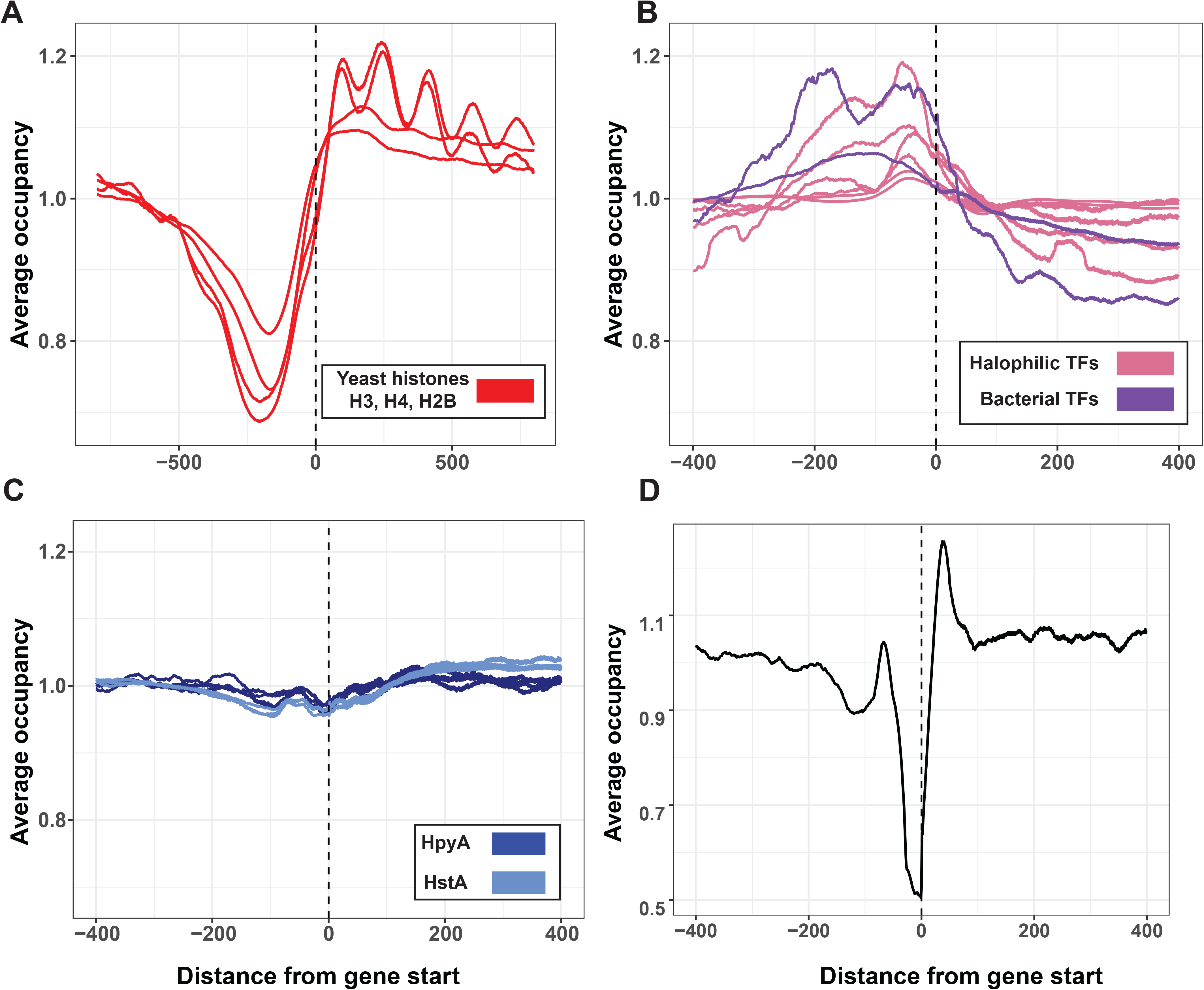
Binding occupancy at start sites of selected DNA-binding proteins. Average binding occupancy across all genes for (**A**) yeast histones (red), (**B**) HpyA (dark blue, 1 representative replicate from each condition tested) and HstA (light blue, 3 replicates, where each line is one replicate), and (**C**) Bacterial TFs *E. coli* FNR and *Brucella abortus* VjbR (purple) and Archaeal TFs *Hca. hispanica* TrmB, *Hfx. volcanii* TroR, *Hfx. med* RosR (pink, 2 replicates each). (**D**) Re-analysis of genome-wide average of micrococcal nuclease digestion (MNase-seq) pattern for *Hfx. volcanii*. In each panel, x-axis represents distance from start site (bp), y-axis represents average occupancy, as measured by read depth in genomic positions around the start site, normalized to average depth across the genome.

In the case of haloarchaeal TFs, we detected, as expected, increased occupancy ~50 bp upstream of gene start sites (**Fig. 4B**), which corresponds with prior data that TFs generally bind in gene promoters(4). Depletion is observed within gene bodies, corresponding to the known relatively infrequent binding within genes(26). The bacterial TF FNR also exhibited an upstream enrichment similar to archaeal TFs, again corresponding to the known preference for bacterial TF binding in promoter regions(65). Bacterial NAPs showed a variety of profiles (**Fig. S4**), including upstream enrichment and/or depletion. In contrast to these other DNA binding proteins, haloarchaeal histone occupancy profiles are relatively flat across the ORF start site, (**Fig 4C**), consistent with our observations that neither HpyA nor HstA binds nearby any particular genomic features (**Fig. 3C**). Importantly, these profiles for HpyA and HstA stand in sharp contrast to those of the other DNA binding proteins compared. Heatmap representations of occupancy data for each individual gene corroborate these findings (**Fig. S5**).

We note that no ChIP-seq data are available in the literature for archaeal histones other than HpyA or HstA. However, genome-wide mapping of nucleosomes [genomic regions resistant to micrococcal nuclease (MNase) digestion] has been carried out using MNase-seq in *T. kodakarensis(42, 46)*, *Methanothermobacter thermoautotrophicus(46)*, and *Hfx. volcanii(66)*. In all these studies, a depletion in occupancy just upstream of the start site, like the nucleosome-free region reported for yeast promoters (and shown using ChIP-seq data in **Fig. 4A**) was reported. We were able to reproduce the same TSS occupancy depletion using the publicly available *Hfx. volcanii* MNase data (**Fig. 4D**); however, this MNase profile contrasts strongly with the HstA ChIP-seq occupancy profile (light blue lines, **Fig. 4C**). By contrast, in the case of yeast histones, the histone ChIP-seq occupancy showing upstream depletion and downstream ordered nucleosomes (**Fig. 4A**) correlates strongly with the known MNase digestion patterns(63). The dissimilarity between *Hfx. volcanii* MNase-seq and HstA ChIP-seq occupancy suggests that HstA is not the chromatin protein for *Hfx volcanii*.

Taken together, these data suggest that the start site occupancy patterns of halophilic histones are unique relative those of canonical eukaryotic or archaeal histones as well as TFs, suggesting a divergent DNA binding function.

### Halophilic genomes lack the dinucleotide periodicity that indicates genome-wide optimization for histone binding

To determine whether halophilic genomes carry a genome-wide 10 bp dinucleotide periodicity signal (GPS) indicative of histone packaging(32, 33, 47), we used power spectrum analysis to detect the GPS of AA/TT/TA dinucleotides in the genome sequences of various archaeal model species (**Fig. 5A**, Methods). The genomes of thermophilic species *Methanothermus fervidus* and *Thermococcus kodakarensis* exhibited a sharp peak in their respective spectral density curves at 10-10.3bp, indicative of periodicity that guides histone binding. This was as expected, as the well-characterised histones of these species are known to wrap DNA and function as chromatin packaging proteins(42, 43). In contrast, periodicity in the same range was not detected for four model halophilic species (*Hbt. salinarum, Hfx. volcanii, Hfx. mediterranei, Haloarcula hispanica*) even though their genomes encode histones. For further comparison, we considered organisms in different branches of the tree of life known to use other proteins besides histones to organize their genomes. *Methanosarcina mazei*(50) and *E. coli* show a periodicity of 10.7-11bp instead of 10bp (**Fig, 5A**). This slightly longer periodic frequency may be indicative of negative supercoiling(67). In contrast, *Sulfolobus solfataricus*, which encodes no histones but uses archaeal-specific proteins such as Alba and Cren7 to package its genome(7), lacks both periodicities.

**Figure 5:**
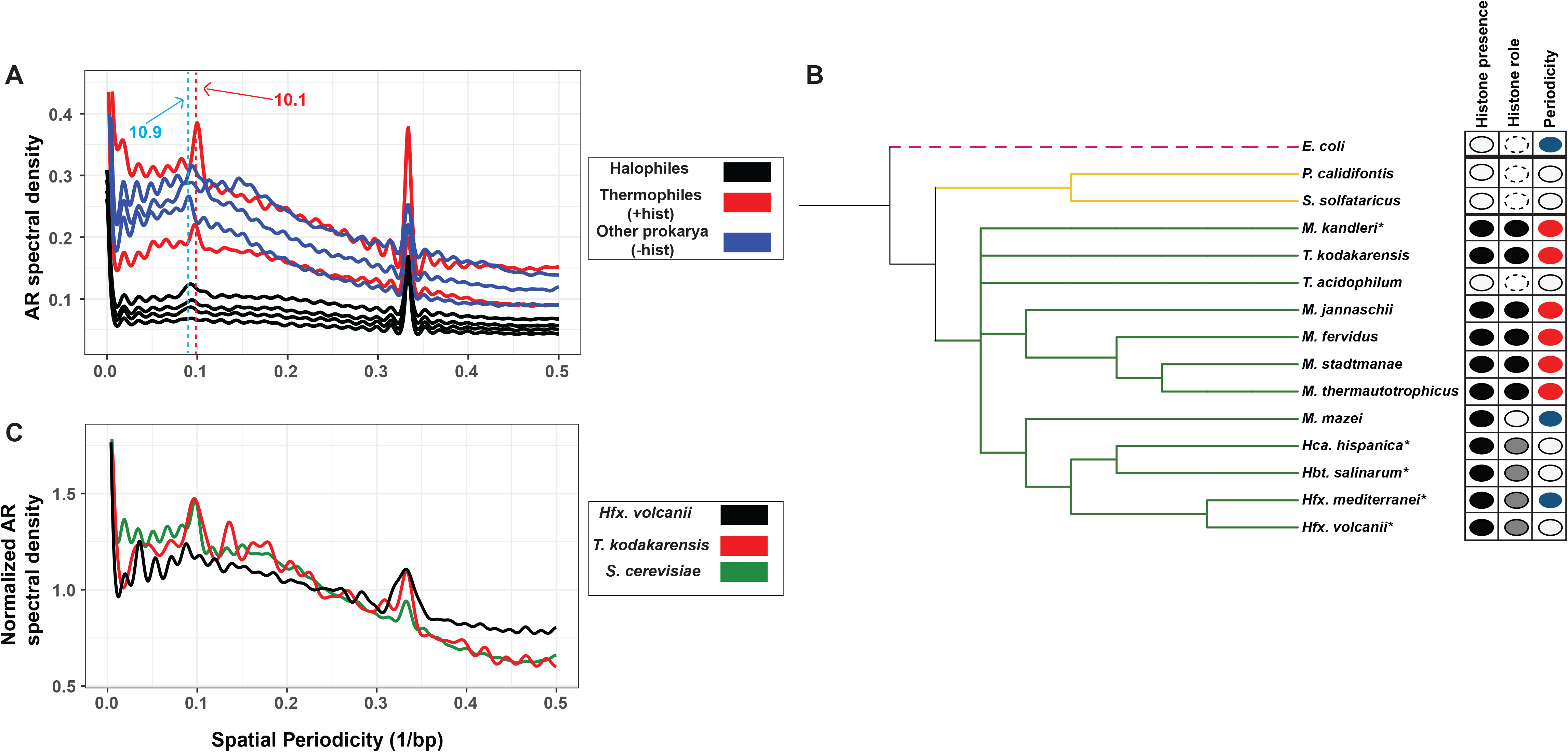
AA/TT/TA dinucleotide periodicity shows histone-linked pattern. (**A**) Autoregression spectra indicating genome-wide dinucleotide periodicity of thermophilic archaeal species with well-characterised histones (*Methanothermus fervidus*, *Thermococcus kodakarensis*; red lines), halophilic archaea that encode histones (*Hbt. salinarum, Hfx. volcanii, Hfx. mediterranei, Hca. hispanica*; black traces) and other prokaryotic species (blue traces) that lack histones (*E. coli*, *S. solfataricus*) or with non-histone chromatin (*M. mazei*). Dotted red line indicates ~10.1bp periodicity present in histone-utilizing species (red traces), dotted blue line represents ~10.9bp periodicity (i.e., from supercoiling) detected in some non-histone utilizing species (blue traces). (**B**) Phylogenetic tree of selected archaeal species (with the bacterium *E. coli* as outgroup). The first column shows detectable (black) or undetected (white) histone-encoding genes. The second column documents experimental characterization of the function of the encoded histone based on previous publications: compaction (black), non-compaction (white), non-canonical histone function (grey), N/A (dotted line). The third column indicates the genome-wide AA/TT/TA dinucleotide periodicity: ~10bp (red), ~11bp (blue), no detectable periodicity (white). Species marked with * indicate high GC content. Only *M. kandleri* genome carries GC periodicity signal. (**C**) MNase protected regions (nucleosomes) of *S. cerevisiae* (green), *T. kodakarensis* (red), *Hfx. volcanii* (black). Dotted black line indicates the ~10bp periodicity peak (not detected for *Hfx. volcanii*). Normalized spectra are plotted in this panel to facilitate clarity in the visualization.

Having validated our method, we extended it first by looking at more archaeal species with experimentally tested and published chromatin proteins. We tested the genomes of *Methanosphaera stadtmanae*, *Methanocaldococcus jannaschii*, *Methanothermobacter themoautotrophicus*, all of which encode experimentally characterised histones(46, 68, 69), as well as *Themoplasma acidophilum* and *Pyrobaculum calidifontis*, which use non-histone chromatin proteins(49, 70). We again detected 10bp GPS in the species using histone to form chromatin but did not detect GPS in those with non-histone chromatin (**Fig. S6A**).

Within halophiles, we also examined the periodicity of the GC dinucleotide, because, unlike the genomes of archaea known to use histones to package their genomes, halophilic genomes are > 60% G+C(71). Therefore, the histone binding signal might be revealed by GC dinucleotides, which are also known to play a role in histone binding(72). *Methanopyrus kandleri*, a GC-rich species(73) whose histone protein has been shown to be capable of nucleosome formation(51), was also included in this comparison. We did not detect a GC dinucleotide 10bp periodicity in halophilic genomes. However, we did observe a periodicity peak very close to 11bp (**Fig. S6B**), which, as noted above, is more likely linked to supercoiling than histone binding. On the other hand, ~10bp GC periodicity was indeed observed for *Mpy. kandleri*.

In summary, across genomes of representative species from Euryarchaea and Crenarchaea, we observe that 10bp GPS is strongly associated with encoded histones that function as chromatin organizers (**Fig, 5B**). In this regard, *Methanosarcina* and halophilic archaea – both with non-essential histones -- are clear outliers.

We next used MNase-seq data from *T. kodakarensis(42)*, *Hfx. volcanii(66)* and the eukaryote *S. cerevisiae(16)* to test if the presence of GPS correlates with MNase-protected regions. These nuclease protected regions have been verified to show nucleosome-based binding periodicity *in vitro* for *S. cerevisiae(16)* and *T. kodakarensis(46)*, but not for *Hfx. volcanii(66)*. We detected a strong 10.2-10.4 bp GPS signal in the protected regions of the two species known to have histone-based chromatin (*T. kodakarensis* and *S. cerevisiae*, **Fig. 5C**). This signal is not detected in the nuclease-protected regions of *Hfx. volcanii*. This evidence combined with the discrepancy between *Hfx. volcanii* occupancy profiles from MNase vs HstA ChIP-seq data (**Fig. 4C, D**) suggests that the histone protein in *Hfx. volcanii* is unlikely to have been the source of previously observed MNase-protected regions(66). This corroborates mass spectrometric proteomics evidence(37, 52) that halophilic histone expression is too low to act as the main chromatin protein in *Hbt. salinarum* and *Hfx. volcanii*. Other potential chromatin proteins in *Hfx. volcanii* are expressed at much higher levels than HstA(74), and are therefore stronger candidates than HstA for generating the MNase-seq TSS depletion pattern (reported previously(66) and reproduced in **Fig. 4D**). mass spectroscopic evidence that Taken together, these data suggest that a genome-wide enrichment for AA/TT/TA periodicity (GPS) of ~10bp is strongly and directly correlated with the presence of histones that function as the main chromatin packaging proteins in archaea. The absence of such GPS (and of 10bp GC periodicity) in halophilic genomes and some archaeal species suggests that their genomic sequence is not optimized for genome-wide histone binding.

### Halophilic histones differ in predicted DNA binding sequence specificity

Because the periodic signal associated with canonical histone binding was absent in halophile genomes, we next asked how HpyA and HstA may bind DNA using *de novo* searches for specific cis-regulatory sequence motifs. Site-specific TFs are usually guided to their target genes by preferential binding to a particular sequence motif(5). This is also true of halophilic TFs, where palindromic motifs have been reported for RosR(75) and TrmB(26) of *Hbt. salinarum*. We employed a variety of *de novo* motif searching methods to detect over-represented cis-regulatory sequences in HpyA- and HstA-bound regions in ChIP-seq data. This included *de novo* motif detection programs such as MEME(76), DNA secondary structures, and over-represented k-mers (Methods; **Supplementary Document S1**).

In the case of HstA, MEME detected (E-value 1.4 x 10^−14^) a palindromic sequence in 31 of the 32 ChIP-seq peaks (**Fig, 6A**; **Supplementary Document S1**). This motif, of the form TCGNSSNCGA (where S is G or C), was robust to correction for background di- and trinucleotide frequencies. Genome pattern scanning analysis using FIMO (part of the MEME suite) detected this motif at 11,630 locations genome-wide, suggesting that HstA may bind additional sites under alternate conditions. In contrast, exhaustive *de novo* computational searches using multiple methods were unable to detect a sequence-specific binding motif for *Hbt. salinarum* HpyA (See details in **Supplementary Document S1**). Instead, we asked whether HpyA binding regions specifically exhibit the periodicity known to facilitate histone binding in other species(33, 46, 47). Although this periodicity was not present at a genome-wide level, a periodicity of 10.4 bp is detected in HpyA-bound regions (**Fig. 6B**). Three of 100 randomly chosen sequences across the genome equal to the length of the HpyA-bound regions exhibited greater spectral peak height (indicating stronger periodicity) in the 10-10.5bp range (**Fig. 6C**), suggesting that HpyA may bind additional sites in the genome and/or under alternative growth conditions not yet investigated. This suggests that, like other histones, HpyA favours binding in DNA regions with a ~10bp dinucleotide periodicity. In contrast, HstA target loci exhibited 11-bp periodicity but not ~10bp periodicity (**Fig. 6D)**. Indeed, the autoregression curve resembles that of the entire *Hfx. volcanii* genome, leaving the cis-regulatory sequence noted above (**Fig. 6A**) as the only determinant of HstA binding detected in our analysis. Taken together, these data suggest that HpyA favours binding to sequences with a ~10bp periodic presence of A/T dinucleotides, implying a histone-like binding mode. In contrast, HstA likely binds in a more sequence-specific manner to a palindromic motif, like TFs.

**Figure 6:**
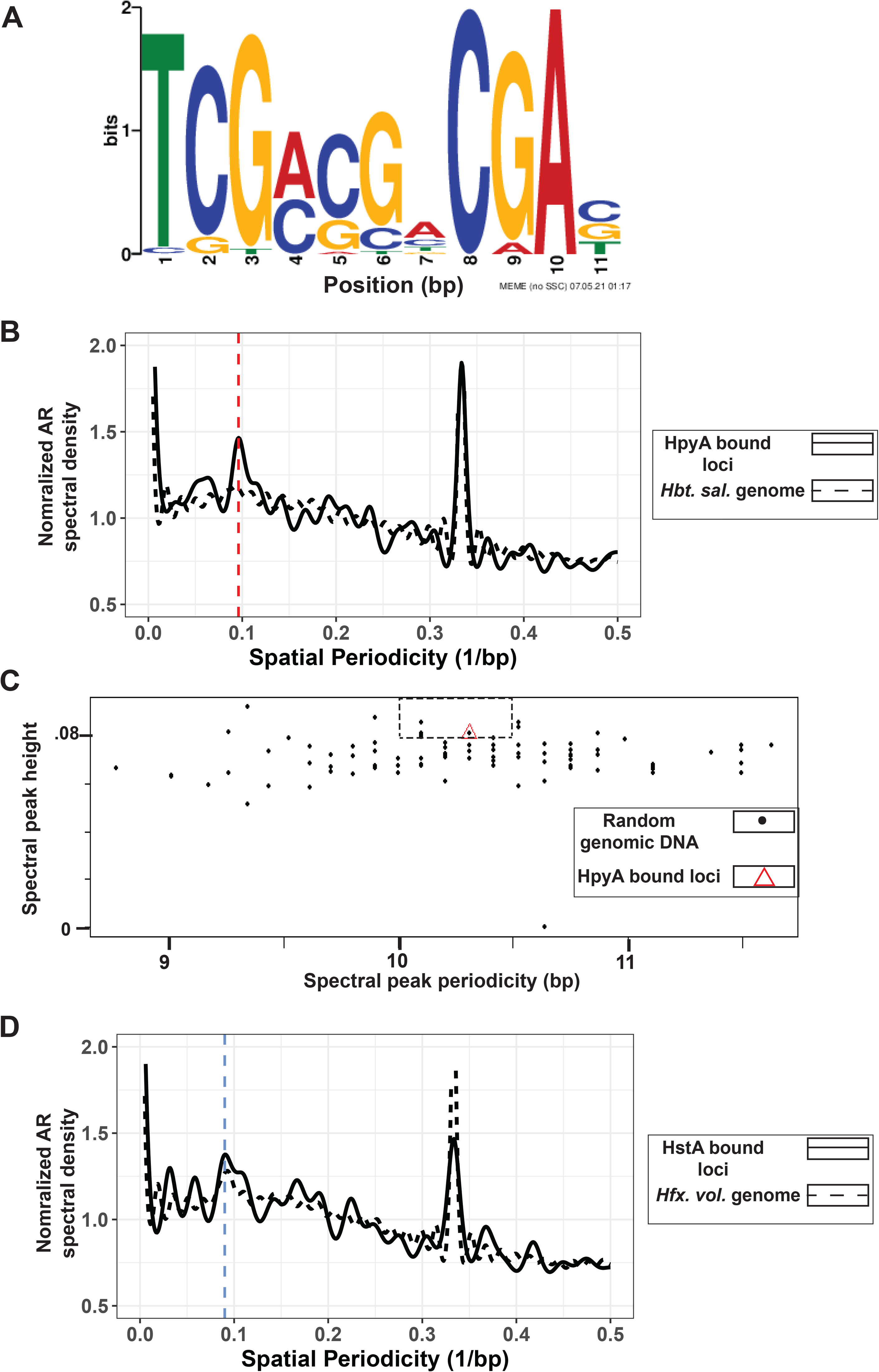
Sequence specificity of HpyA and HstA binding. (**A**) Motif logo of cis-regulatory sequence detected in HstA-bound sites. Bit scores are shown in the y-axis and bp positions on the x-axis. Motif logo generated by the MEME suite(76) output. (**B**) 10.4bp periodicity is present in HpyA-bound loci (solid black line) but absent in the *Hbt. salinarum* genome (black dotted line). Vertical red dotted line indicates 10bp. (**C**) Comparing randomly chosen regions of the genome (black dots) with the periodicity of the HpyA-bound loci (red triangle). Dotted rectangle includes those randomly chosen sequences that show stronger periodicity than HpyA at relevant levels (10-10.5bp). (**D**) ~11bp frequency of HstA-bound loci (black solid line) matches periodicity of the entire genome of *Hfx. volcanii* (black dotted line). Blue dotted line indicates 11bp.

## CONCLUSIONS

Here we demonstrate functional conservation of histone-like proteins across related species of halophiles, with some subtle differences. The sole histone coding gene of two halophilic species is non-essential for growth (**Fig. 1**).However, unlike HpyA of *Hbt. salinarum*, HstA of *Hfx. volcanii* is important for growth under optimum conditions. The genome-wide binding of HstA and HpyA are similar to one another with respect to their pattern of binding: number, width, and shape of peaks; percentage of genome covered; and lack of preference for genomic features. Interestingly, HpyA and HstA differ in terms of their binding site sequence preferences. In the case of *Hbt. salinarum*, HpyA-bound regions were observed to contain 10bp dinucleotide periodicity. HstA, on the other hand, may bind a TF-like palindromic sequence motif. In comparing halophilic histones to DNA binding proteins across domains of life, we observe a pastiche of conserved features. Considering this evidence, we conclude that HpyA and HstA functions diverged from those of other archaeal and eukaryotic histones. This is consistent with and extends knowledge from our previous characterization of HpyA in *Hbt. salinarum*, where we demonstrated its function as a transcriptional regulator of ion uptake(56). Given the strong sequence and structural conservation of histones across sequenced halophile genomes(52), we posit that alternative functions are likely for all halophilic histones. Our results are consistent with the hypothesis that archaeal histone function varies according to habitat through the process of selection under extreme conditions(37).

Chromatin proteins are highly expressed(24, 37), and ChIP-seq data demonstrates that their genome-wide binding results in a large number of peaks covering over 10% of the genome (**Fig. 3B**, see also refs(20, 21)) or in characteristic occupancy signals in the case of eukaryotic histones (**Fig. 4A**, see also ref.(63)). Previous proteomics mass spectrometry results from our lab(52) and others(37) demonstrated a low expression level of HpyA in *Hbt. salinarum* and HstA in *Hfx. volcanii*, comparable to that of a TF, and therefore too low to provide architectural organization of the genome. The sparse binding patterns of HpyA and HstA corroborate this hypothesis (**Fig. 2**). More broadly across the archaeal phylogenetic spectrum, histone expression level is strongly associated with chromatinization of the genome(37). Hence, HpyA and HstA differ from chromatin proteins with respect to the binding frequency necessary for architectural functions. Taken together, these data suggest that the DNA binding patterns of halophilic histones are unique relative those of canonical eukaryotic or archaeal histones as well as TFs, suggesting a divergent DNA binding function.

These results therefore situate halophilic histone-like proteins in a growing group of DNA binding proteins that defy categorization according to the criteria commonly used to differentiate between them (**Fig. 7)**. Dorman and colleagues posited that traditional definitions of bacterial DNA binding proteins as “TFs” or “NAPs” are insufficient to capture the true continuum of functional characteristics observed for certain proteins(5). For example, some proteins were defined as NAPs because of their ability to bind genome-wide and alter DNA structure; however, some NAPs can also bind in a highly sequence-specific manner (e.g., IHF). Some TFs like CRP exert sequence-specific control of certain loci but bind to hundreds of other sites in the genome(77, 78). Such examples are not restricted to bacteria: newly discovered site-specific archaeal TFs are also likely to bend or loop DNA. Examples include the TetR family TF FadR(79, 80) and archaeal Lrp family proteins(28, 81). Depending on the locus, Lrp family proteins can bind with or without sequence specificity(23), exhibit direct or indirect effects of transcription(25), combining features observed for TFs as well as chromatin-like proteins. The DNA binding proteins under investigation in the current study clearly require more flexible functional categorization. We conclude that halophilic histones, with their primary sequence homology to archaeal and eukaryotic histones(38, 52), their role as transcription regulators(52, 56), and hybrid modes of DNA binding, lie within the unclear divide between TFs, histones, and nucleoid associated proteins.

**Figure 7:**
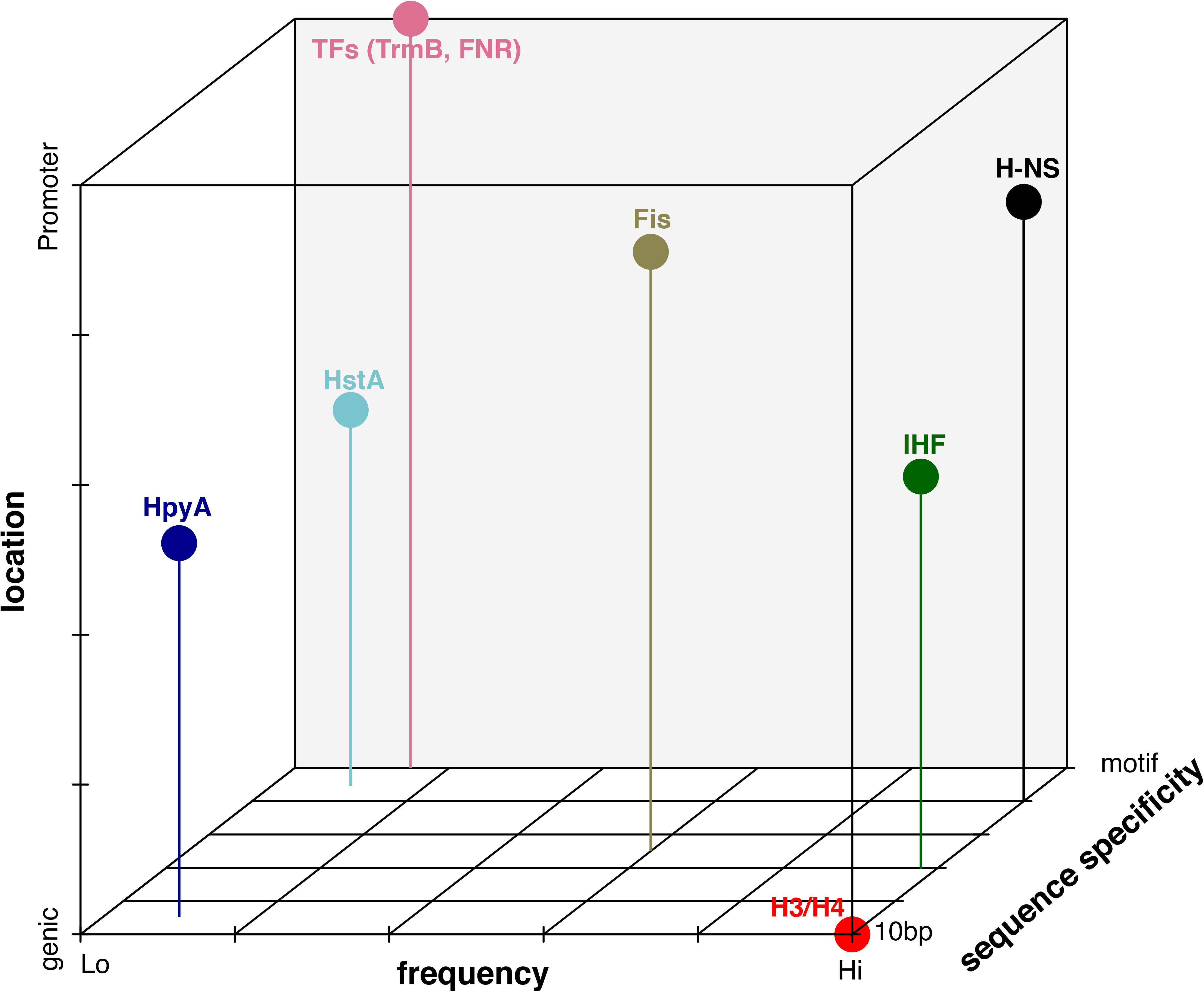
Visual summary of binding characteristics of selected DNA-binding proteins investigated in this study. Binding is classified based on sequence specificity, ranging from preference for 10bp periodicity (10bp) to strict *cis* sequence motif (motif); frequency as measured by genome-wide coverage and number of peaks, ranging from low (Lo) to high (Hi); location preference, ranging from coding to promoter preference. Figure is qualitative: tick marks and gridlines are intended for visual clarity.

## MATERIALS AND METHODS

### Strain construction

The *Haloferax volcanii* wild type strain used in this study was DS2(82). The strains created here used DS2 derivative Δ*pyrE* (strain H26) as the parent strain. The Δ*hstA* (*HVO_0520*) knockout strain AKS198 was created from parent H26 using vectors described by Allers *et al.(83)* and the pop-in pop-out double crossover counterselection strategy commonly used for *Hfx. volcanii(83)*. Briefly, the pAKS145 knockout vector was generated by isothermal ligation of sequences flanking the *hstA* gene into backbone vector pTA131 at the EcoRV site. Strains, primers, and plasmids used for all strain constructions are noted in **Table S6**.

AKS214 was the strain used to test in *trans* complementation of Δ*hstA* deletion growth defect. It contains the pAKS147 plasmid, which was created by inserting *hstA* and 500bp of its upstream sequence into the pJAM809 backbone at the XbaI and KpnI sites. Two strains were generated for ChIP-seq experiments. AKS217, the negative control strain, is the Δ*hstA* background carrying the pJAM809 empty vector(84). AKS233 is the Δ*hstA* strain carrying plasmid pAKS180, which was derived from pAKS147 by addition of the hemagglutinin (HA) tag using the NEB Q5 site-directed mutagenesis kit.

*hstA* deletion from the genome and *hstA* or *hstA-HA* presence in-*trans* was confirmed with PCR and Sanger sequencing of the flanking regions. Deletion was additionally confirmed with full-genome resequencing. Full-genome resequencing for the parent strain and Δ*hstA* strain was analysed using the *breseq*(85) analysis tool, results are given in **Table S6**.

### Media, culturing, and phenotyping

*Hfx. volcanii* rich medium was used for routine growth across experiments: Yeast Peptone Casamino Acids (Hv-YPC), as described previously(83). For plasmid maintenance, media were supplemented with Novobiocin (0.1 μg/mL). For construction of deletion mutants, media were supplemented with 5-FOA (300μg/mL) in selection of the second crossover.

To measure growth rates of Δ*pyrE* parent strain and Δ*hstA* strains, at least 3 biological replicate individual colonies of H26 and AKS198 were picked from plates freshly streaked from frozen stock and precultured for 70-80 hrs in 5mL Hv-YPC at 42°C with 225 rpm shaking (referred to as “standard” or “optimum” conditions” throughout). To test growth phenotypes in standard conditions, precultures were diluted to OD_600_~0.025 and then cultured in a BioScreen C (Growth Curves USA) at 42°C with fast shaking at maximum amplitude. Each biological replicate culture was inoculated into at least duplicate and up to quadruplicate wells of the microtiter plate to ensure technical reproducibility in the measurements. OD600 was measured by the BioScreen every 30 minutes over the growth curve. Further details of stress conditions tested and results, are given in **Fig. S1**. Resultant growth curves were quantified by measuring the area under the log-transformed growth curve (AUC) Visualization and area under the curve (AUC) analysis of the growth curve was carried out as in https://github.com/amyschmid/Halophilic_histone_binding/tree/main/Growth_analysis. Growth data for Δ*hstA* and parent strain in optimal conditions is provided in **Table S1**.

### ChIP-seq experiment

*Haloferax volcanii* HstA-HA ChIP-seq was carried out using the same method as described previously(56). Briefly, three biological replicate cultures of AKS233 (*hstA*-HA) and 1 replicate of AKS217 as a negative control were grown in 50 mL of YPC18% and harvested at 15-17 hours post-inoculation at an optical density of 0.21-0.33 (mid-exponential phase). Cultures were cross-linked, immunoprecipitated by virtue of the HA tag, and DNA prepared as described previously. Strain details are provided in **Table S6**. As before, the Duke Center for Genomic and Computational Biology carried out library preparation including adapter ligation. The only difference from previous protocol was use of the Illumina NovaSeq6000 to carry out paired-end sequencing.

### ChIP-seq analysis

Publicly available ChIP-seq data from the relevant TF, NAP, or histone was downloaded from the NCBI sequence Read Archive using the fastq-dump feature from SRAToolkit 2.9.0 (https://hpc.nih.gov/apps/sratoolkit.html). Details of published datasets used for bacterial and archaeal TFs, histones, and NAPs, including sequence read archive trace numbers, are provided in **Table S5**. The bacterial NAP HU was excluded from this analysis because it does not show IP/Input enrichment, indicative of binding, like other TFs/NAPs do(21). Fastq files were converted to sorted BAM files, wig files, and per-base read-depth text files were generated as described previously(56).

Sorted BAM files were used to generate peak lists using the R package MOSAiCS(86). Peak lists were created for haloarchaeal TFs (*Hca. hispanica* RosR and TrmB, *Hfx volcanii* RosR and TroR, and *Hfx mediterranei* RosR), selected bacterial NAPs (IHF, H-NS, FIS), and bacterial TF (FNR). The bacterial TF VjbR was excluded from this analysis because the vast majority of the peaks called by Mosaics could not be visually confirmed. For experiments where more than one replicate from the same conditions was present, multiIntersectBed from Bedtools(87) was used to combine peaks across replicates, and only peaks present in the majority of replicates were considered. The code used to generate this is provided in https://github.com/amyschmid/Halophilic_histone_binding/tree/main/Peak_calling.

The peak list for HpyA was taken from our previously published work(56). The peak list for HstA was created as described above, but with some manual curation (details below). The average width and total area covered by the peaks within these lists was calculated within Excel, and total area covered was expressed as a percentage of genome length (**Table S3**).

Peaks were classified as “intergenic” or “coding” based on where in the genome they were located. The centre of each peak was found and was determined to be within or outside a coding region (as described by the list of genes in the NCBI gene table for that species). The code used to make this classification, and to graph the results, is in https://github.com/amyschmid/Halophilic_histone_binding/tree/main/Bindingfeatures.

The results of this classification were used as the basis of a hypergeometric test in R using the phyper function: phyper (#peaks in non-coding regions, length of genome that is non-coding, length of coding genome, #total peaks) to determine if peaks were over-represented in intergenic regions (**Table S4**).

### Generating *Hfx volcanii* HstA peak list

As mentioned above, MOSAiCS was used to generate peak lists from HstA ChIP-Seq data, and peaks common in at least 2 of 3 replicates were retained to make a joint peak list. This list was then curated manually to remove false positives caused by changes in input control sequencing, transposase and integrase-caused local duplications, and peaks common with the HA tag-alone input control. The final manually curated peak list for HstA is noted in **Table S2**.

### Start site occupancy analysis

Per-base read-depth text files were generated as described above. These text files were used as inputs for occupancy analysis, alongside genome annotations downloaded from NCBI (details in **Table S5**). Code was written that returns a matrix where each row corresponds to a single gene, and the columns represent sequence depth at positions from ±400 bp of that start site, normalized to the average depth over the whole chromosome. (For yeast analysis, considering the larger size of intergenic regions, and the known length of DNA bound to a single nucleosome as 147bp, this analysis was repeated by changing the boundaries to ±800bp). The occupancy graph was generated by taking the average of occupancy across all start sites (rows in the matrix). The code used for this analysis is in https://github.com/amyschmid/Halophilic_histone_binding/tree/main/TSSgraphs.

Note that the start site used here refers to the ORF translation start site instead of the transcription start site (TSS) that is often used for these analyses. Three criteria motivated this choice: (a) ORF start sites are better annotated in most species including halophilic archaea; (b) ORF and transcription start sites are very close in halophiles, with >60% of ORFs being leaderless in *Hfx. volcanii*(88); (c) using ORF start sites, we were able to reproduce previously seen patterns for yeast TSS(63) (**Fig. 4A**).

### Dinucleotide periodicity analysis

FASTA files containing the genome sequence of the relevant species were downloaded from the NCBI website (species and download details in **Table S5**) and were analysed using custom R scripts. In brief, dinucleotides (AA/TT/TA) are detected in each genome and binarized: locations with these dinucleotides are marked as 1 and the rest of the genome as 0. Then, the autoregression spectrum spec.ar() function in R (with default parameters) was used to estimate the spectral density of this binary signal, which indicates the periodicity of the selected dinucleotides using an autoregression fit. For facilitating clarity in visualization of autoregression curves, periodicity was normalized by the average signal in **Figs 5B, 5C**, and **S5**. The same analysis was carried out for GC dinucleotides to generate **Fig. S5B**. For nucleosome enrichment analysis, data regarding the centre of the nucleosomes was downloaded from supplementary information of Brogaard et al 2012(16) (for *Saccharomyces cerevisiae*), Maruyama et al 2013(42) (for *Thermococcus kodakarensis*), and Ammar et al 2012(66) (for *Haloferax volcanii*). Sequences of the length of a typical nucleosome (150bp for eukaryotes, 30-60bp for archaea) were isolated around each centre and the same analysis as above was carried out. Note that the strong peak at 0.33bp^−1^ (3bp) seen in all these spectra is linked to codon usage; it is present in all species and is not linked to histone binding(89). Depending on the AT content of the sequence being examined, some of the spectra have an increasing or decreasing slope resulting from slight deviations in A+T content locally; this too is not linked to histone binding(89). The codes used for these analyses are available at https://github.com/amyschmid/Halophilic_histone_binding/tree/main/Periodicity_genomewide.

The results of the genome-wide periodicity obtained and shown in **Figs 5A** and **S5** are summarized in **Fig. 5B**. The phylogenetic tree for this figure was made using the Integrated Tree of Life (iTOL, https://itol.embl.de).

### Motif search

Bed files containing peak locations from HstA and HpyA ChIP-seq data were converted to FASTA format using the Bedtools(87) getfasta command. These FASTA files were used as input for various motif and over-represented sequence determining programs. We used motif-detection with MEME(76) and Homer(90), k-mer detection tool KMAC(91), and a DNA secondary structure detection R-package called gquad (https://cran.rproject.org/web/packages/gquad/index.html). Finally, the fasta-get-markov tool of MEME was used to determine background mono-, di- and tri-nucleotide frequencies. A more detailed description of the parameters used for each program, and the results of the searches, is provided in **Supplementary Document S1**.

For obtaining periodicity of sequences bound by HpyA or HpyA, the FASTA file containing all the peaks (generated as above) was merged into a single line. Finally, periodicity of the sequences in this FASTA file were analysed. A similar method to the procedure used to analyze genome-wide periodicity was used here. The obtained periodicity was compared with randomly chosen sequences from the genome roughly equal in length to the width of ChIP-seq peaks (peak widths given in **Supplementary Table S2** and in reference(56)). These simulated peak lists were then analysed using the autoregression spectrum scripts, and results compared between the 100 simulated sequences and the empirically detected peaks.

## DATA AVAILABILITY

The ChIP-seq data have been deposited in the National Center for Biotechnology Information (NCBI) Gene Expression Omnibus (GEO) accession number GSE186415. The whole genome sequencing data for the Δ*hstA* deletion strain have been deposited in the NCBI Sequence Read Archive at accession PRJNA773760. All code and input data for analyses presented here are freely available via the GitHub repository https://github.com/amyschmid/Halophilic_histone_binding. All supplementary figures, tables, and documents are available on FigShare at https://doi.org/10.6084/m9.figshare.19391648.v1.

## CONTRIBUTIONS

S.S. and A.K.S devised the project and wrote the manuscript. A.K.S secured the funding for the project and contributed to data analysis. S.S. carried out strain creation, HstA ChIP-seq, and data analysis under supervision of A.K.S.

C.D., A.H., M.M.P. and R.H. carried out ChIP-seq of halophilic transcription factors, provided technical assistance and expertise for experiments and analysis. M.M.P and R.H. edited the manuscript.

## ACKNOWLEDGEMENTS

The authors thank all Schmid lab members for their feedback on the study and comments on the manuscript. S.S. acknowledges the support of his graduate thesis committee for their comments and advice on the study (David McAlpine, Amy Grunden, Richard Brennan). Funding for the study was provided by grants MCB-1651117 and 1936024 from National Science Foundation to AKS.

## Supplementary material

*All supplementary material are available via FigShare at https://doi.org/10.6084/m9.figshare.19391648.v1*

### Document S1

Details of methods and results obtained when searching for a DNA sequence motif for HstA and HpyA binding.

**Figure S1: Δ*hstA* phenotype in response to diverse stress conditions.** (**A**) Raw growth curves of multiple biological replicate cultures under standard growth conditions (YPC medium, 2.5M NaCl, 0.3 M Mg^2+^, 42°C). (**B**-**E**) Each graph depicts the area under the log-transformed growth curve (AUC) under the conditions indicated at the top of each panel. Each point represents AUC for one growth curve, and the horizontal lines depict the median of the AUC distribution for each strain under each growth condition. (**B**) Growth in increased sodium (4M NaCl); (**C**) growth in reduced sodium (1.5M NaCl). (**D**) gluconeogenic conditions (no sugar added to HvCa medium). HvCa medium contains identical components to YPC18 described above, except without the addition of peptone or yeast extract(83). (**E**) Oxidative stress (0.05M H_2_O_2_). Δ*hstA*: parent growth ratio in all cases was 0.84 or higher, indicating *hstA* is not required for stress response. (**F**) Growth curves in high MgCl_2_ (overall Mg^2+^ 0.48M), low MgCl_2_ (overall Mg^2+^ 0.19M); visual analysis of the data confirmed that growth under Mg stress conditions was identical to non-stress conditions.

**Figure S2: HstA deletion can be complemented *in trans*; HA tag does not interfere with HstA function.** Dot plot representing growth measured by area under the curve (AUC) of Δ*hstA* containing plasmids as calculated by the R package grofit; each dot represents AUC for one technical replicate growth curve, central line represents median value. Samples from left to right: *ΔhstA* expressing wild type *hstA in trans; ΔhstA* expressing wild type *hstA* fused to the HA tag; Δ*hstA* strain transformed with empty vector control.

**Figure S3: HpyA and HstA bind in few, discrete peaks, like TFs, and contrasting with histones and NAPs.** (**A**) HpyA (dark blue) and HstA (light blue) binding is observed as discrete reproducible peaks (marked with arrow); shown here are 2 representative replicates each. (**B**) Halophilic TF TrmB (pink) binds in discrete reproducible peaks; 2 replicates shown (**C**) Bacterial TF FNR (purple) binds in discrete reproducible peaks (2 replicates shown). (**D**) Bacterial NAPs cover a large part of the genome, either with many peaks: IHF (green), FIS (olive); and/or broad peaks: HNS (black). (**E**) Yeast histone ChIP-seq (red) shows depletions in promoter regions of some genes (representative regions marked with black arrows). Broad peak locations of nucleosomes positioned in the gene body are indicated with grey dots. Shown here are data from histone H2B, H4, H3 in quiescent phase (H3_Q), and H3 in logarithmic growth phase (H3_log).

**Figure S4: Binding occupancy at start sites of bacterial NAPs.** Average binding occupancy across all genes for *E. coli* IHF (green, 2 subunits), H-NS (black), FIS (olive).

**Figure S5: Heatmap of binding occupancy for yeast histone and halophilic histone.** (**A**) Heatmap of yeast histone H3, with each row representing the start site of one gene. Colour scale at right represents average normalized occupancy. (**B**) Line graph of histone H3 showing the average across all genes. (**C**) HpyA heatmap with colour scale as in A. (**D**) HpyA average line graph.

**Figure S6:** (**A**) AA/TT/TA dinucleotide periodicity of archaeal species with characterised histones (red; *M. stadtmanae, M. thermoautotrophicus, M. jannaschii*) and with non-histone chromatin (blue; *T. acidophilum, P. calidifontis*). Dotted red line represents 10 bp periodicity peak location. (**B**) Periodicity of GC dinucleotides in species with >60%G+C genomic content: histone-containing *M. kandleri* (red), model halophile species *Hbt. salinarum, Hfx. volcanii, Hfx. mediterranei, Hca. hispanica* (black). Red dotted line indicates 10bp, black dotted line indicates 11bp.

**Table S1:** Two tabs in the supplementary excel file: (a) Growth data (OD600) for Δ*hstA* and Δ*pyrE* parent strain in optimal conditions; (b) whole genome resequencing data for Δ*hstA* strain.

**Table S2:** Manually curated list of ChIP-seq peaks for HstA.

**Table S3:** Details of characteristics (number of peaks, average width, total area covered by peaks) of ChIP-seq peaks for HpyA, HstA, and other DNA-binding proteins (shown graphically in main text figures).

**Table S4:** Results of hypergeometric test to check over-representation of ChIP-seq peaks in intergenic regions of the genome.

**Table S5:** List of ChIP-seq datasets used (with SRA trace where available), and genomes analysed for periodicity (with link to the relevant NCBI assembly).

**Table S6:** List of primers, strains, and plasmids used in this study.

